# 3D brain angiogenesis model to reconstitute maturation of functional human blood-brain barrier *in vitro*

**DOI:** 10.1101/471334

**Authors:** Somin Lee, Minhwan Chung, Noo Li Jeon

## Abstract

Human central nervous system (CNS) vasculature in brain expresses a distinctive barrier phenotype, the blood–brain barrier (BBB), which protects the brain against harmful pathogens. Since the BBB contributes to low success rate in CNS pharmacotherapy by restricting drug transportation, the development of an *in vitro* human BBB model has been in demand. Previous models were unable to fully represent the complex threedimensional (3D) anatomical structure or specific barrier phenotypes of the matured BBB. In this study, we present a physiological 3D microfluidic model of the human BBB that mimics its developmental process including CNS angiogenesis and subsequent maturation in concert with perivascular cells. We used microfluidic hydrogel patterning to precisely and sequentially load perivascular cells into the model, investigate the role of each cell type on BBB phenotypes. We confirmed the necessity of the tri-culture system (brain endothelium with pericytes and astrocytes) to attain the characteristic BBB vascular morphology such as minimized diameter and maximized junction expression. In addition, endothelial-perivascular cell interaction was also critical in reconstituting p-glycoprotein (p-gp), efflux transporter in our model that works as metabolic barrier of BBB and blocks drug to enter CNS. The 3D hydrogel matrix was tuned with hyaluronic acid (HA) to optimize the interaction between endothelial cells and astrocytes. Our *in vitro* BBB system mimics CNS angiogenesis and characteristic features of BBB. We expect the model will contribute to deeper understanding of neurodegenerative diseases and cost-efficient development of effective CNS medications.

## 1. Introduction

Blood vessels in the brain cooperate with neuronal and perivascular cells to develop a highly specific microenvironmental niche called the neurovascular unit. The endothelium within the neurovascular unit forms the blood–brain barrier (BBB), which exhibits stronger and more selective barrier function than that of other tissues. The barrier function of the BBB is important to maintaining homeostasis in the CNS and protects it from pathogens; however, it also blocks numerous drug molecules that may useful for treating CNS diseases [1, 2].

Specifically, highly developed junction proteins between endothelial cells (ECs) physically restrict hydrophilic compounds, such as polar drugs, from entering the CNS while allowing the passage of essential hydrophobic molecules including oxygen and hormones. The transport of small hydrophilic molecules is tightly regulated by specific transport systems on luminal and abluminal membranes of brain ECs in an ATP-mediated manner. Nutrients are selectively permitted to enter the CNS, whereas potentially harmful materials are blocked or actively released from cells by specialized transporters [3]. Importantly, these unique BBB functionalities are acquired via complex cellular interactions during developmental angiogenesis of the CNS [4–6]. Although some molecular mechanisms involved in BBB phenotypes have been discovered, the complex interplay between BBB cell types has yet to be fully elucidated. Thus, much effort has been devoted to developing *in vitro* models to mimic BBB physiology since many genes of interest related to BBB have shown embryonic lethality [4].

As cellular interactions among brain ECs, astrocytes, and pericytes in the BBB microenvironment are critical to developing the BBB phenotype, *in vitro* models have emphasized the importance of co-culture. Static models using a transwell culture system were first introduced in the 2000s; these studies focused on measuring transendothelial electrical resistance (TEER) to determine how pericytes and astrocytes contribute to junction development and barrier integrity. Although early transwell models used mouse brain ECs [7, 8], Hatherell et al. [9] developed a tri-culture transwell model using immortalized human ECs, pericytes, and astrocytes. The strength of these models was the simplicity of their culture methods, however, they could not recreate the dynamic fluid conditions of the CNS vascular niche or the complex physiological anatomy of the BBB.

Recently, microfluidics-based models have made progress towards developing *in vitro* BBB models by providing surmounting these limitations. Ross et al. [10] presented the first microfluidic BBB model, which was designed with a monolayer of mouse EC cells over a porous membrane co-cultured with or without mouse astrocyte cells. This system was optimized to measure TEER for the analysis of endothelial permeability under various conditions. The model introduced fluidic shear stress, which was impossible to attain in previous static models. A number of studies inspired by this model have employed microfluidic BBB models to study BBB functionality and drug delivery [11–15]. These models demonstrated poor physiological resemblance to the human BBB due to absence of human primary cells, and failed to accurately represent the 3D structure of the luminal vasculature within the surrounding 3D extracellular matrix (ECM) microenvironment.

Within the past 3 years, several 3D BBB vascular models have appeared. Cho et al. [16] designed a 3D BBB model to study neurovascular pathology in a rat brain EC cell line and introduced neutrophil, which extravasates from the 3D lumen in the presence of a tumor necrosis factor-alpha (TNF-α) chemotactic gradient. Herland et al. [17] used human-derived primary brain ECs, astrocytes, and pericytes in a 3D model to show natural interactions among cells in the absence of any artificial barrier, such as a porous membrane. *In vitro* models of the neovascular unit consisting of 3D vasculature with neurons and astrocytes have also been introduced and contributed to the establishment of a co-culture protocol for ECs and neuronal cells [18, 19]. However, among these models, 3D vasculature is recreated artificially by attaching ECs to the walls of the pre-lumenized ECM or microfluidic channel. This protocol overlooks the process of CNS vasculature development, which is critical to the acquisition of unique phenotypes, such as narrow vascular morphology and high efflux transporter expression in BBB.

Exceptionally, CNS vasculature is not developed via vasculogenesis which is canonical mechanism for common vasculature development. Rather than vasculogenesis, angiogenesis is the main mechanism for development of CNS vasculature in which ECs sprout from the perineural vascular plexus to invade the embryonic neuroectoderm towards a gradient of a proangiogenic factor such as vascular endothelial growth factor (VEGF-A) [4, 20]. Several review articles have explained that BBB development occurs in three sequential steps: angiogenesis, differentiation, and maturation [2, 4, 20]. EC phenotypes change in each phase; for example, transcytotic vesicles and leukocyte adhesions are robust in the early angiogenesis phase and decrease during the maturation phase. In contrast, the expression of tight junctions and efflux transporters is increased between the differentiation and maturation phases, and some CNS angiogenesis-specific molecular systems, such as Glut-1, are regulated by Wnt-β-catenin signaling, which is related to barrier-specific properties [2, 21]. Moreover, Umans et al. also reported that barriergenesis of BBB occurs simultaneously with CNS angiogenesis *in vivo* using zebrafish mutant model [6]. These examples clearly show the strong interconnection between the maturation of BBB functionality and CNS angiogenesis (Fig. 1B).

**Figure 1.**
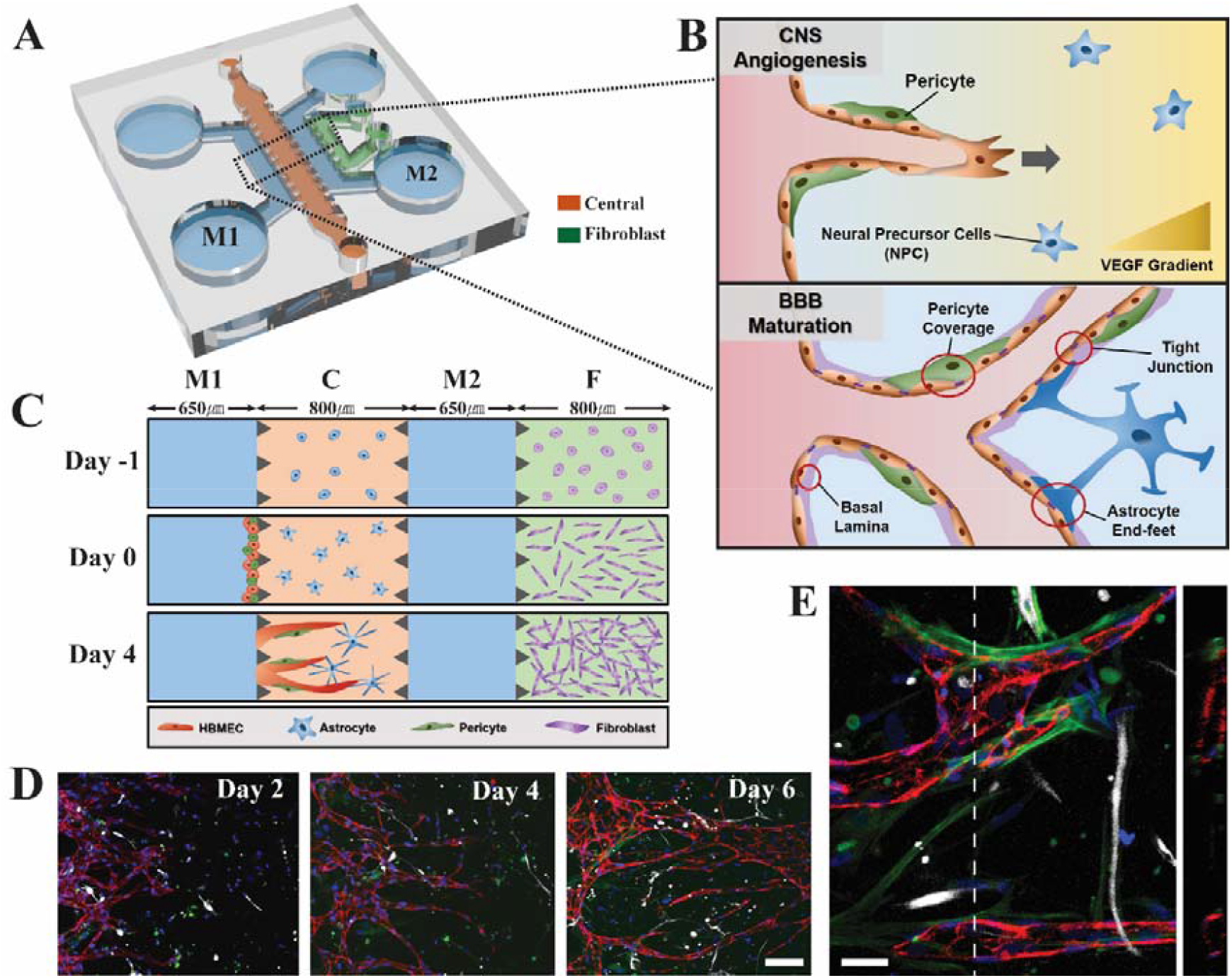
Design and cell culture configurations of microfluidic platform mimicking CNS angiogenesis. (A) Schematic view of microfluidic device having three types of microfluidic channel. (B) Biological concept of the model to reconstruct 3D *in vitro* BBB by mimicking CNS angiogenesis which sequentially maturated the BBB phenotypes. (C) Section view of microfluidic device describing sequential loading and culture progress of BBB. Channel C (red) is the region where final BBB microenvironment is constructed. Channel F (green) possess 3D fibroblast which acts as source of angiogenic factors. Channel M1 and M2 (blue) are media channel. (D) Day by day confocal imaging of channel C where HBMEC (anti-CD31, red) sprouts from left end to right end. Astrocytes (anti-GFAP, white) protrude to generate end-feet and pericytes (anti-αSMA, green) wrap around EC as day goes by. Nucleus were stained by Hoechst 33342 (blue). Scale bar: 100μm. (E) Confocal imagining at day 6 in higher magnification. Cells were labeled same as in (D). Right image is the section view of white dotted line in left image, describing that the vessels are lumenized in 3D. Scale bar: 30μm. 3D reconstruction of the same image is presented in supplementary video 1.

We have established a microfluidic-based *in vitro* model by mimicking *in vivo* CNS angiogenesis, thus maximizing the physiological relevance of the model to BBB development. We designed the model to reconstitute the 3D CNS microenvironment, wherein BBB functionality is developed. In this model, all three types of primary human cells (brain ECs, astrocytes, and pericytes) have an *in vivo*-like 3D morphology, with direct cellular interactions occurring within the microfluidic channel. By mimicking the sequential process of BBB development, we were able to confer *in vivo*-like morphology and barrier function on our *in vitro* model. We focused on verifying the p-gp efflux transporter system, which is representative of the BBB metabolic barrier mechanism. This functionalized *in vitro* system will potentially serve as a novel platform to reveal unknown human BBB biology and test the efficiency of drug delivery systems across the human BBB.

## 2. Materials and Methods

### 2.1. Microfluidic device fabrication

Microfluidic device was fabricated by polydimethylsiloxane (PDMS, Slygard 184, Dow Corning) using replica molding. The master mold was made through photolithography on silicon wafer using photoresist, SU-8 100 (MicroChem) resulting patterns of microstructure having height of 150μm. Liquid sate of PDMS prepolymer was completely mixed in 10:1 (w/w) ratio of PDMS base and curing agent, then poured on the master to be polymerized for 30 minutes on hot plate. PDMS device was gently peeled off from the master, then media reservoirs and hydrogel inlets/outlets were punched using biopsy punch and sharpened hypodermic needle (18 gauge), respectively. Finally, the device was cleaned with adhesive tape to remove dusts or extra PDMS particles, then plasma bonded to be covalently bonded with clean coverslip. As plasma bonding made the device hydrophilic inadequate for patterning hydrogel channels, it was stored on 70°C dry oven for at least two days before loading hydrogel in order to restore its hydrophobicity.

### 2.2. Cell culture

Primary human brain microvascular endothelial cells (HBMEC, CellSystems) were cultured in endothelial basal medium-2 (EBM-2, Lonza) supplemented with EGM-2 MV BulletKit (Lonza). Primary human umbilical vein endothelial cells (HUVEC, Lonza) were cultured in EBM-2 supplemented with EGM-2 BulletKit (Lonza). Both ECs were cultured in dish to have less than 80% of confluency during passaging or experiments. In addition, they are both used in experiments at passage 4 or 5. Primary human pericytes from placenta (hPC-PL, PromoCell) were cultured in pericyte growth medium (PGM, PromoCell) and used in experiments at passage 7 or 8. Primary normal human astrocytes (NHA, Lonza) were cultured in astrocyte growth medium (AGM, Lonza) and used in experiments at passage 6. Primary normal human lung fibroblasts (LF, Lonza) were cultured in fibroblast growth medium (FGM, Lonza) and used in experiments at passage 6. All cells were maintained in culture condition of 5% CO_2_ and 37°C inside humidified incubator.

### 2.3. Cell seeding for CNS angiogenesis

For our 3D cell culture of BBB microvasculature, fibrin hydrogel was basically used for ECM. The fibrin hydrogel solution (10mg/ml) was prepared by fibrinogen from bovine plasma (Sigma-Aldrich, F8630) diluted in phosphate-buffered saline (PBS, Hyclone), then mixed with aprotinin solution (0.45U/ml, Sigma-Aldrich, A1153) in volume ratio of 25:4. To generate 3D cellular matrix inside the microchannel, cell suspension is mixed with prepared fibrin hydrogel solution at volume ratio of 3:1 to yield final fibrin concentration being 2.5mg/ml. The cellular hydrogel solution is injected inside microfluidic channel as soon as it is quickly mixed with thrombin solution (1U/ml, Sigma-Aldrich, T4648) in volume ratio of 50:1, resulting cross-link of hydrogel around in 3 minutes.

In order to generate BBB microenvironment for CNS angiogenesis, different types of cells were sequentially loaded inside the microchannel over two days. All of media used in cell culture on microfluidic BBB platform is a mixture of EGM2-MV and AGM in 1:1 ratio, in order to maximize viability and physiological characteristics of both ECs and astrocytes. In first day, LFs were mixed with fibrin hydrogel solution and loaded on the side channel F in final cell concentration of 6×10^6^ cells/ml. Astrocytes followed after being 3D patterned by mixing with fibrin hydrogel solution and loaded on the center channel C in final concentration of 4.5×10^6^ cells/ml. After 5 minutes later while hydrogel was crosslinked enough, media was introduced on top two media reservoirs among four total reservoirs. As the chip is initially hydrophobic during hydrogel loading step, the media could be entirely fill inside the media channels, channel M1 and M2, by vacuum aspiration from other two unfilled media reservoirs. Media was filled enough and samples were incubated at 37°C overnight. Next day, all media in four reservoirs were aspirated while slight amount of media remain inside the media channels. The device was tilted in 90° as channel M1 and M2 goes up and down, respectively. Then, 5μℓ of cell suspension, mixture of HBMECs and pericytes in ratio of 6:1 (total cell concentration is 5×10^6^ cells/ml), was injected on one of the reservoirs connected to channel M1 in order to make cells to be attached on the left side of previously loaded 3D hydrogel matrix in channel C. The sample was incubated at 37°C as it was tilted for 30 minutes which is enough for cells to form focal adhesions on fibrin ECM matrix. Finally, all reservoirs are filled with medium and cultured in humidified incubator at 37°C and 5% CO_2_.

The sequential cell loading progress described above is one of four co-culture conditions conducted in this research, condition EAP having all composing BBB cells in a 3D microenvironment. For condition E and EP, channel C is filled with acellular fibrin hydrogel (final concentration is 2.5mg/ml) at the first day. For condition E and EA, cell suspension being introduced to be attached on the side of the central channel C at the second day is only composed by ECs (final concentration is 5×10^6^ cells/ml). To avoid the effect of medium component dominating the effect of cellular interactions, culture media are same in all co-culture conditions.

### 2.4. Immunostaining

All samples were fixed with 4% paraformaldehyde (Biosesang) for 15 minutes. In advance to introducing antibodies, samples were pre-treated with 0.2% triton-X 100 (Sigma-Aldrich) during 20 minutes for permeabilization. Treatment of 3% bovine serum albumin (BSA, Sigma-Aldrich) followed after for 1 hour in order to avoid unspecific bonding of antibodies. All chemical solutions were treated at room temperature and they are introduced inside the microchannel by adding around 30μℓ on top two reservoirs after aspirating remaining liquid on four media reservoirs. After BSA treatment, the samples were filled with PBS and stored in 4 before immunostaining.

Antibodies as a marker for specific cell type were introduced after removing PBS. Alexa Flour 488/594/647-conjugated mouse anti-human CD31 (1:200, Biolegend; 303110/303126/303112) was used as EC marker. Alexa Fluor 488-conjugated mouse anti-human GFAP (1:200, BD Bioscience; 560297) was used as astrocyte marker and Alexa Fluor 594-conjugated mouse anti-human α-SMA (1:200, R&D systems; IC1420T) was uses as pericyte marker. They are all treated for 2 days. Hoechst 33342 (1:1000, Invitrogen; C10339) was used for nuclear staining done in an hour. For endothelial junction staining, Alexa Fluor 594-conjucated mouse antihuman ZO-1 (1:200, Life Technologies; 339194), Alexa Fluor 488-conjucated mouse anti-human Claudin5 (1:200, Invitrogen; 352588) and Alexa Fluor 488-conjugted mouse anti-human VE-Cadherin (1:200, eBioscience; 53-1448-42) were used and treated for 2 days. For basement membrane staining, unconjugated primary antibodies such as rabbit anti-human laminin (1:100, Abcam; ab11575) and rabbit anti-human collagen IV (1:100, Abcam; ab6586) were treated for 2 days, then Alexa Fluor 568 goat anti-rabbit IgG (1:1000, Invitrogen; A11036) was treated as secondary antibody for overnight. All of samples were preserved in 4°C during immunostaining and maintained in PBS until imaging. For p-glycoprotein staining, rabbit anti-human pgp (1:100, Abcam; ab129450) was used for 2 days and aforementioned secondary antibody was treated for overnight.

### 2.5. Imaging and image analysis on vessel morphology

Confocal laser scanning unit (Olympus FV1000) with IX81 inverted microscope (Olympus) was used to reconstruct 3D microenvironment of BBB network. Images were captured by confocal PMT detector and x10, x20 lenses were used (Olympus). Confocal images were processed by software called IMARIS (Bitplane).

Live cell imaging for calcein-AM efflux test was performed using inverted fluorescence microscope (Nikon Eclipse Ti, Japan) and microscope software NIS elements was used for multi-stage and time-lapse imaging (Nikon). Images in this microscope was captures using Hamamastsu Orca R2 camera at 16-bit depth and x20 lens was used to analyze individual vessels in vasculature network (Nikon).

Fiji (http://fiji.sc.), open access software, was used to analyze confocal images of BBB vasculature. Z-projection of whole stack could result misreading of branch information since two or more vessels will overlap in same position, the sample stacks were divided in lower and upper parts before the projection. After original 3D images were projected and converted into 2D binary masked images, we then conducted measurement of vessel morphology. Total vessel area was calculated in ratio (%) of total vascular area value over total matrix area value ranging from start point of branching. Starting point of branching in each co-culture condition varied so the vessel area was analyzed in ratio of coverage. Average vessel diameter was measured along a line that vertically divides the vessel network into half. All pixel values, 0 or 255 since they were binary images, along the line were measured and we determined the vessel boundary as the value changes to 255. We measured the length of line having continuous value of 255 and recorded as vessel diameter in each sample.

### 2.6. Western blot assay

Expression level of endothelial cell adhesion molecule CD31 (PECAM-1) and endothelial tight junction protein ZO-1 were quantified by western blot assay. PDMS microfluidic chips for the assay were exceptionally attached on PSA film without plasma bonding, instead of cover glass with plasma bonding. This fabrication enabled detachment of chips after culture. After 7 days of culture after EC attachment, samples were washed with PBS for three times. PSA film was removed gently and cellular hydrogel remaining on PDMS piece was lysed in RIPA buffer (T&I) with 1x protease inhibitors. Lysed samples were collected and total protein concentration of each sample was quantified by Bradford assay. 15μg of protein was separated on SDS-PAGE gel (8%) and transferred to PVDF membrane (GE Healthcare). 5% skim milk in TBS-T buffer (25mM Tris, 190mM NaCl and 0.05% Tween 20, pH 7.5) was used to block PVDF membrane for an hour, then primary antibodies, rabbit anti-ZO-1 antibody(250:1, Abcam), rabbit anti-CD31 antibody (250:1, Abcam) or rabbit anti-β actin (1000:1, Abcam) was applied on each membrane overnight at 4°C. Samples were washed again for three times, 30 minutes each, by TBS-T buffer. IgG-HRP secondary antibody was introduced in concentration of 500:1 or 1000:1 for 1h 30minutes at room temperature. Visualization was performed using the Clarity Western ECL substrate (Bio-Rad). Band intensities were measured using Fiji software.

### 2.7. P-glycoprotein inhibitor test by Calcein-AM efflux assay

Samples cultured for 7 days after EC attachment were pretreated with Valsopodar (Adooq), also known as PSC-833, in concentration of 10μM for 10 hours. Control groups were also pretreated with DMSO added media at the same time. Then, media of all four reservoirs were removed in each chip and clacein-AM (ThermoFisher) diluted in concentration of 2μM was added on top two reservoirs, 150μℓ individually. After 30 minutes, solution was removed and inhibitory media and DMSO control media was added on reservoirs again for live imaging. Multiple regions of each condition were selected and total 59 regions were captured every 20 minutes by using multi-stage and time-lapse imaging program. We finally gained 33 timeframes for each region for 640 minutes of imaging. For each time frame image, we specifically designated five squared points (Fig. 4D, yellow boxes in 0 hour images) in same size which covers one endothelial cell consisting the lumenized vessel. The fluorescence intensity of each squared point was measured for all 33 frames and we analyzed the change of intensity over time. Moreover, change in intensity (ΔI=I_t=0 min_ - I_t=640 min_) over initial intensity (I_t=0_) was measured (ΔI/I_t=0_) in each squared point in every sample.

### 2.8. Fibrin-HA ECM matrix preparation

Hyaluronic acid (HA) was prepared by HyStem™ Cell Culture Scaffold Kit (Sigma) which consist of HyStem and Extralink1 and they were both prepared in concentration of 10mg/ml. HA and crosslinker was mixed in volume ratio of 4:1. Then the mixture was finally added on prepared fibrin hydrogel solution with cell suspension right before loading in the microfluidic channel. Final concentration of fibrin was fixed as 2.5mg/ml, on the other hand, HA’s final concentration was controlled as 0mg/ml, 1.5mg/ml and 2.5mg/ml in each condition.

## 3. Results

### 3.1. The Microfluidic model mimicked CNS angiogenesis and reconstituted the BBB microenvironment in a 3D ECM

To reconstitute the human BBB system, we used our previously developed microfluidic platform [22] which has four channels compartmentalized by arrays of micro-posts (Fig. 1A and 1B). Briefly, channel C is the main region of interest, wherein the 3D BBB microenvironment is developed. Channel M1 and M2 are positioned next to channel C and connected to reservoirs for media supply. Channel F is positioned next to channel M2 to pattern the fibrin hydrogel-embedded 3D stromal cell culture. Channels C and F were filled with human astrocytes and human normal LFs, respectively, and stabilized in 3D fibrin hydrogel for 1 day. HBMECs were then mixed with human pericytes and attached along the M1-C boundary (Fig. 1C). Brain ECs exhibited sprouting from the left side to the right side of channel C, and pro-angiogenic factor gradients were generated by LFs in channel F. Interestingly, pericytes and astrocytes exhibited physical associations with newly formed 3D brain microvessels. After 6 days of culture, ECs reached the other side of the channel and the blood vessel network was perfused (Fig. 1D). We confirmed the perfusion and integrity of the vascular network with FITC-dextran (70 kDa) (Supplementary Fig. 1A). Most parts of the network were perfused regardless of vessel diameter or density, making the platform eligible for various types of transluminal assays. Importantly, pericyte coverage was observed along microvessel surfaces, whereas multiple astrocyte end-feet directly contacted the perfused network, resembling the constitution of the BBB microenvironment (Fig. 1E, Supplementary Video 1).

### 3.2. Endothelial-perivascular cell interaction regulates shape of the vascular network generated by CNS angiogenesis

It is widely acknowledged that interactions between brain ECs, perivascular cells, and their corresponding microenvironments determine the distinct character of the BBB [3, 23–29]. However, these complex interactions have yet to be simulated by current *in vitro* BBB models.

To investigate how co-cultured perivascular cell types interact with blood vessel cells in our system, we first analyzed the morphology of the blood vessel network co-cultured with endothelial astrocytes (EAs), endothelial pericytes (EPs), or both cell types (EAPs) (Fig. 2A). Perivascular cells suppressed blood vessel dilation. The total area and branch width of the vessels significantly decreased under co-culture. Interestingly, EA- and EP-suppressive effects were synergetic under tri-culture (EAP).

**Figure 2.**
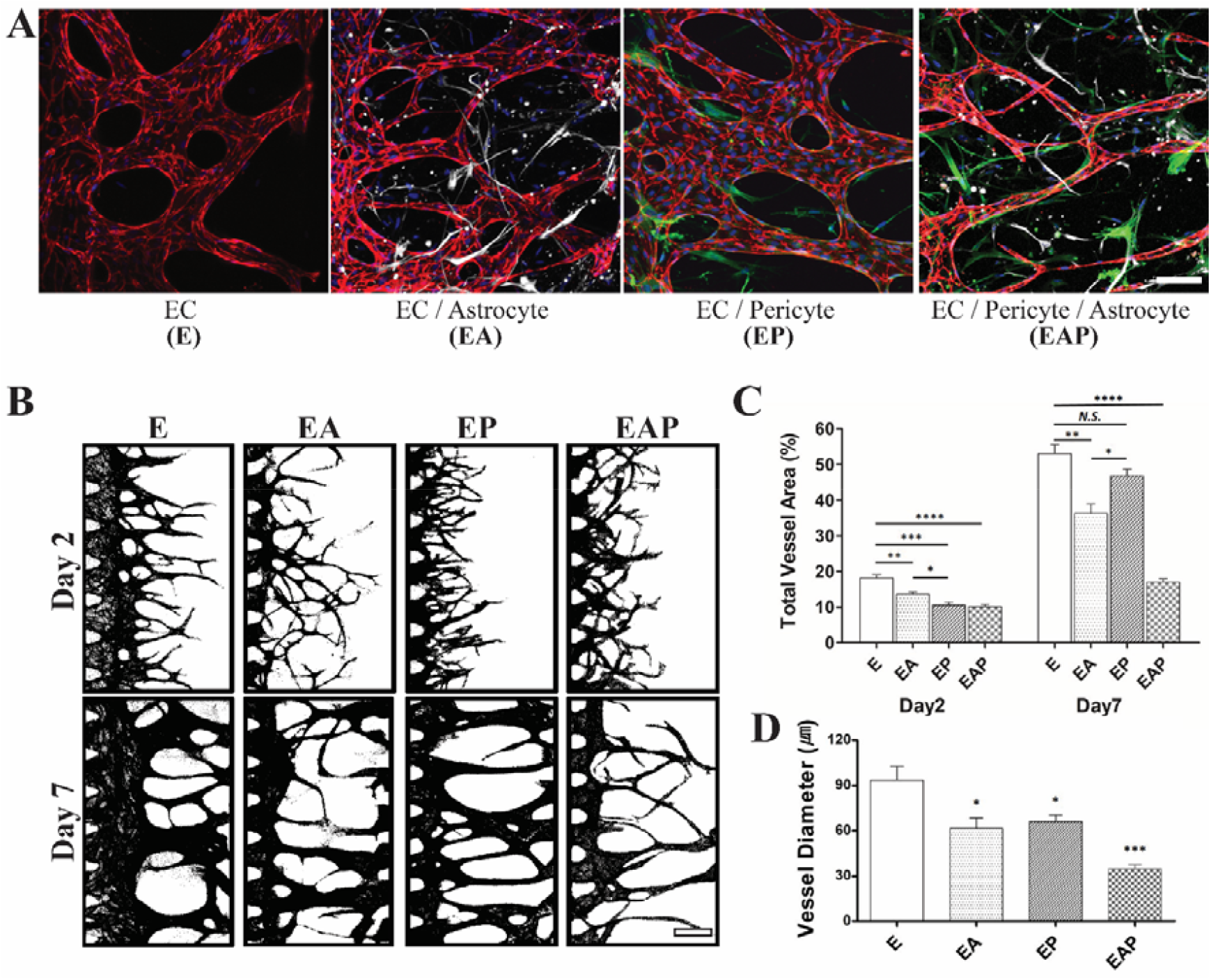
Analysis of vascular morphology regulated by different types of perivascular cells. (A) Confocal images of 3D vasculature in each co-culture condition. HBMEC, pericytes, astrocytes and nucleus were labeled by anti-CD31(red), anti-αSMA (green), anti-GFAP (white) and Hoechst 33342 (blue), respectively. Scale bar: 100μm. (B) Masked images of 3D vasculature in each co-culture condition fixed at day2 and day7. Scale bar: 200μm. (C) Quantitative analysis on total vasculature area at day 2 and day 7 in each co-culture condition. (D) Quantitative analysis on average vessel diameter at day 7 in each co-culture condition. (C, D) N=8 in day 2 condition EA, EP, day 7 condition E, EA and EP; n=10 in day 2 condition E, EP, day 7 condition EAP. Error bars represent SEM. Unpaired two-tailed Student’s t-test was performed to obtain statistical comparisons of analyzed values, with the *p* value threshold for statistical significance set at *p<0.05; **p<0.005; ***p<0.0005; ****p<0.00001.

The ratio of total vessel area to total ECM matrix area in channel C was measured for each sample (Fig. 2C). We selected this analysis protocol instead of analyzing actual area values because the starting points of branches from the left end of channel C differed among co-culture conditions, as illustrated in the masking images of vessel network (Fig. 2B). Condition E resulted in 53.02% of coverage at day 7, which was the maximum among all conditions. Under condition EA at day 7, some branches were of similar width to those under condition E, but others were much narrower. Vessels were sparsely distributed along the overall network under condition EA, resulting in the second lowest vessel area percentage (39.27%). Under condition EP, the vascular network covered 46.71% of the total vessel area at day 7. Under tri-culture, the BBB network had significantly lower coverage under condition EAP, at 17.01%. Differences in vascular geometry among co-culture conditions were not significant during the early stage of CNS angiogenesis, according to the total vessel area at day 2 post-EC loading. This result indicates that morphological characteristics were gained through cellular interactions between ECs and perivascular cells throughout the angiogenic progress. We also examined vessel diameter (Fig. 2D). The lowest average vessel diameter was observed under condition EAP at day 7, at 34.64μm, which was 2.7-fold narrower than that under condition E and 1.9-fold narrower than that under conditions EA and EP.

We conducted the same co-culture experiment using human umbilical vein endothelial cells (HUVECs) instead of HBMEC; the origin of these cells does not match the CNS microenvironment. Vessel morphology results for each co-culture condition were very different from those of the HBMEC experiments (Supplementary Fig. 2A). We confirmed this finding by quantitative analysis of the total vasculature area at day 5 (Supplementary Fig. 2B). Unlike HBMECs, HUVECs under the tri-culture condition did not yield the smallest vascular areas. The largest vessel coverage was observed in the presence of astrocytes and ECs only. These results indicate the importance of the origin of ECs to the development of *in vitro* BBB models.

### 3.3. Expression of BBB specific junction proteins

A distinguishing phenotypic feature of maturation during BBB development is the significant expression of junction proteins and basal lamina proteins on ECs. Therefore, we verified these phenotypes in the model by immunocytochemistry and western blot analysis. The expression of endothelial tight junction proteins ZO-1 (Fig. 3A), Claudin-5 (Fig. 3C), and adherens junction protein VE-cadherin (Fig. 3C) in HBMECs was individually visualized by confocal microscopy.

**Figure 3.**
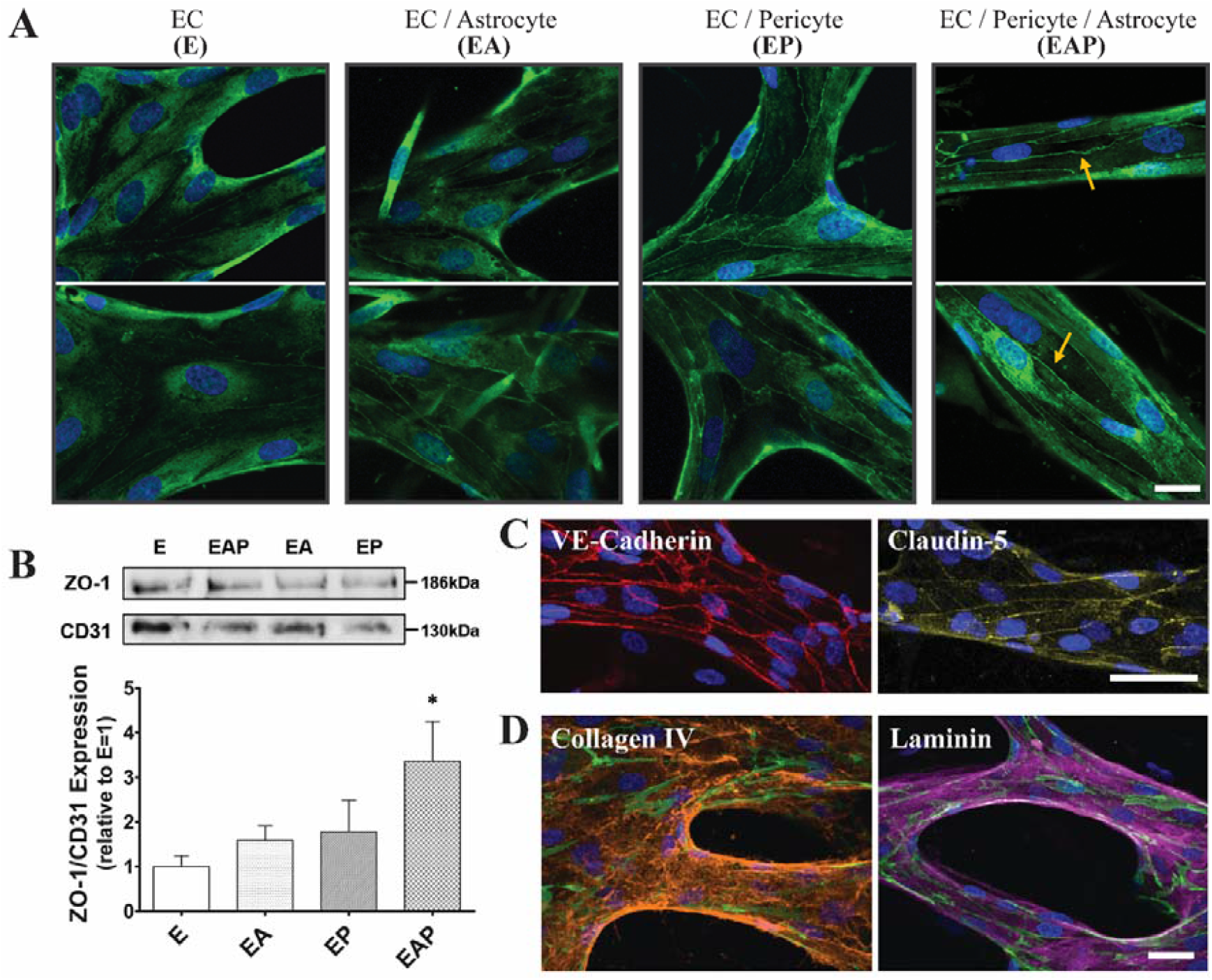
Analysis on expression of junction and basal lamina protein in each co-culture condition. (A) 3D confocal images of ZO-1 tight junction expression at day 7 culture labeled with anti-ZO-1 (green) and Hoechst 33342 (blue). Two representative images were displayed for each co-culture condition. ZO-1 expression on cellular boundary in condition EAP is pointed with yellow arrows. Scale bar: 20 μm. (B) Western blot assay for quantifying ZO-1 expression. Upper image is representative band image of ZO-1 and CD31 for each co-culture condition. Below graph is the result of quantitative analysis on ZO-1 expression level in each co-culture condition normalized against the expression level of CD31. N=6 for condition E, EA and EAP; n=5 for condition EP. Error bars represent SEM. Unpaired two-tailed Student’s t-test was performed to obtain statistical comparisons of analyzed values, with the *p* value threshold for statistical significance set at *p<0.05. (C) 3D confocal image of adherens junction VE-Cadherin (red) and tight junction Claudin-5 (yellow) in tri-cultured vasculature at day 7. Nucleus were labeled by Hoeschst 33342 (blue). Scale bar: 50μm. (D) 3D confocal image of basal lamina proteins, Collagen (orange) and Laminin (purple), in tri-cultured vasculature at day 7. ECs and Nucleus were labeled by anti-CD31 (green) and Hoeschst 33342 (blue), respectively. Scale bar: 30μm.

We were able to clearly confirm differences in expression intensity and geometry due to co-culture conditions, particularly for ZO-1 expression. Four representative images of anti-ZO-1 immunostained (green) HBMECs under different co-culture conditions were taken at the same settings (Fig. 3A). Under condition E (HBEMCs mono-culture condition), ZO-1 expression was not significant around cell-to-cell boundary, instead distributed over the cell body. When HBEMCs sprouted in the presence of astrocytes (condition EA), the boundaries between ECs were visible through the ZO-1 junction. However, the expression signal was weaker than that under conditions EP and EAP. Condition EAP exhibited the strongest ZO-1 intensity and a clear zipper-like boundary between ECs. These differences in fluorescence intensity indicate that the expression of the ZO-1 tight junction protein was determined by cellular interactions between ECs and surrounding BBB perivascular cells. Moreover, the highest expression level in presence of all types of BBB cells show that BBB maturation was reconstituted in the model.

We confirmed the immunofluorescence result with western blotting (Fig.3B). Since the number of ECs in each condition varied (Supplementary Fig. 3A), ZO-1 expression level was normalized to the amount of endothelial cells in each sample by CD31 expression level. The ratio of average expression level of ZO-1 over CD31 reached a maximum under condition EAP (2.458) which was 3.36-fold higher than that of condition E (0.731). Ratio of expression level in condition EA (1.162) and condition EP (1.562) was both higher than that of condition E. This quantitative result was consistent with the result of the immunofluorescence intensity analysis.

The expression of basal lamina proteins, such as collagen IV and laminin, was then verified in the 3D BBB *in vitro* model. Under condition EAP, vasculature was immunostained after being fully matured for 7 days, and exhibited deposition of basal lamina around the perivascular surface. Confocal images clearly showed a layer of ECM proteins marked with either anti-collagen IV (orange) or anti-laminin (purple), and wrapping ECs marked by anti-CD31 (green) (Fig. 3D).

### 3.4. Functional efflux pump eligible for multi-drug resistant assay

Efflux transporters on the surfaces of brain ECs prevent intracellular accumulation of the substrate and exhibit primary BBB barrier function. These transporters restrict the entry of drugs, resulting low success rate of CNS drug discovery [29, 30]. Among these transporters, p-gp is a well understood ATP-driven efflux pump that is multi-specific to various drugs [30–32]. Therefore, p-gp inhibitors were highlighted for use in drug co-treatment, resulting in enhanced effectiveness of the drugs [30].

Thus, we first determined p-gp expression by immunofluorescence imaging in tri-cultured *in vitro* BBB microvessels. In section view, p-gp (red) was spatially located in the intraluminal side of the ECs (Fig. 4A, yellow arrowhead). To demonstrate the usability of our co-culture system as a drug testing platform, we tested the functionality of efflux transporters in our model under different co-culture conditions. Based on previously described calcein-AM fluorometric functional assays [33, 34], we used calcein-AM molecules as a substrate of p-gp to trace the transcellular dynamics of CNS drugs (Fig. 4B). A fully perfused vascular network of EC monoculture or BBB tri-culture was pre-treated with the p-gp inhibitor Valsopodar (PSC-833; Adooq) for 10 hours at a concentration of 10μM. Calcein-AM (C3100MP; Thermo Fisher) was then introduced at a concentration of 2μM to be uptaken by ECs. The efflux was monitored every 20 minutes for total 640 minutes using a live imaging system (Fig. 4C).

**Figure 4.**
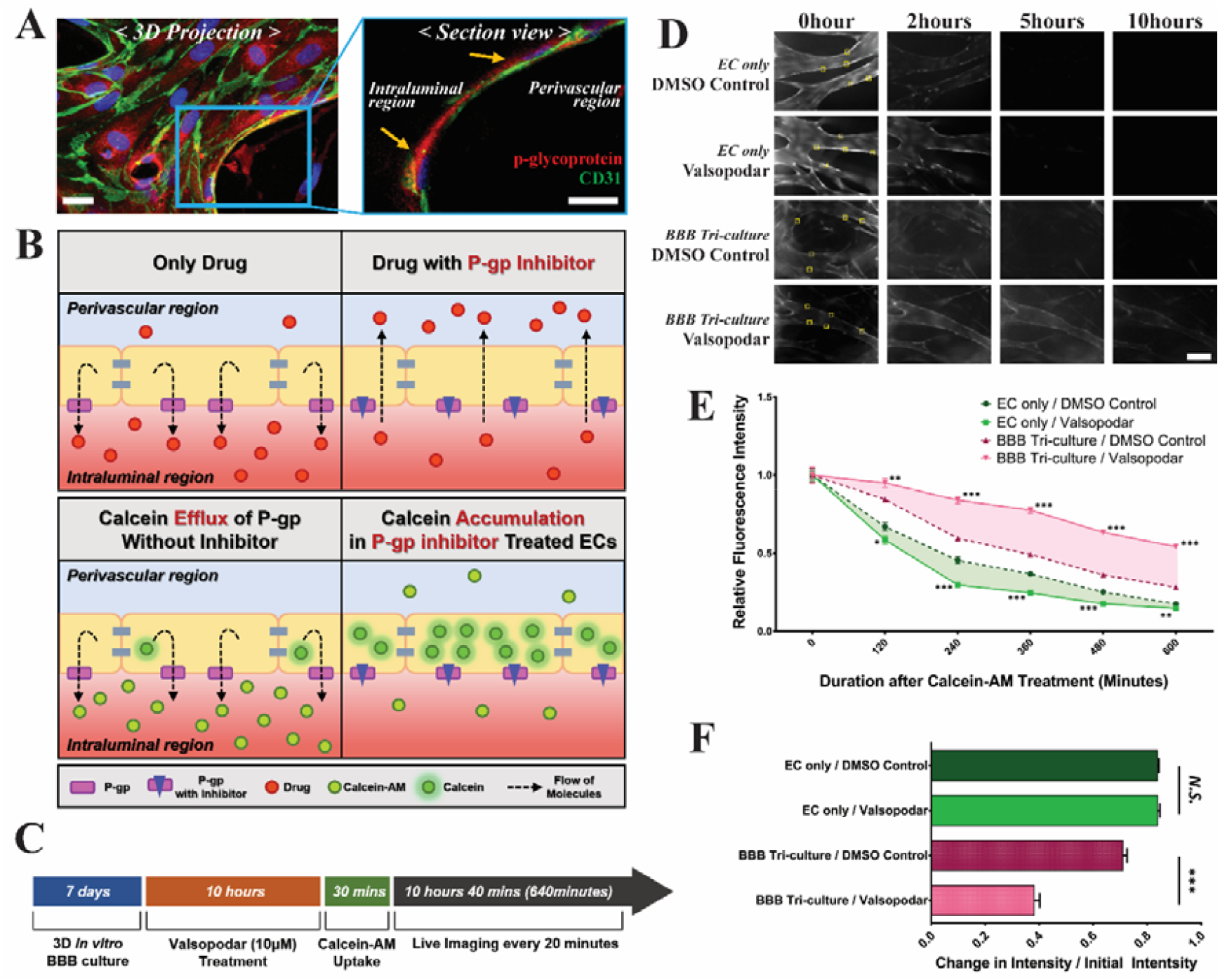
Verification of functional efflux pump in developed *in vitro* BBB model by testing p-gp inhibitor via calcein-AM efflux assay. (A) 3D confocal image of p-gp protein (red) expressed in tri-cultured vasculature at day 7. Right section view is zoomed from left 3D projection image which describes intraluminal oriented expression of p-gp (pointed as yellow arrow heads). ECs and Nucleus were labeled by anti-CD31 (green) and Hoeschst 33342 (blue), respectively. Scale bar: 20μm. (B) Conceptual description of calcein-AM efflux assay (bottom) which mimics actual effect of p-gp inhibitor on drug resistance by p-gp *in vivo* (top). (C) Schematic description of work flow for calcein-AM efflux assay in developed *in vitro* BBB with or without p-gp inhibitor Valsopodar. (D) Representative image of calcein fluorescence in each time point. Vasculature was cultured in EC mono-culture or BBB tri-culture condition with or without p-gp inhibitor pre-treatment. Scale bar: 100μm. (E) Change of calcein fluorescence intensity at the designated region on vessel wall (yellow boxes in Fig. 4D 0hour images) every 120 minutes. (F) Quantitative comparison of change in calcein fluorescence intensity during total experiment duration (640 minutes) normalized by initial intensity. (E, F) n=65 for EC only/DMSO Control; n=70 for EC only/Valsopodar treated; n=50 for BBB Tri-culture/DMSO control; n=110 for BBB Tri-culture/Valsopodar treated. Error bars represent SEM. Statistical comparisons between analyzed values of DMSO control and Valsopodar treated group in each time point was obtained from unpaired two-tailed Student’s t-test, with the *p* value threshold for statistical significance set at *p<0.05; **p<0.005; ***p<0.00001.

Intracellular fluorescence diminished over time (Fig. 4D and 4E), however, the rate of decrease differed between culture conditions. Interestingly, whether the inhibitor was pre-treated or not influenced the efflux rate of intracellular calcein only under BBB tri-culture. ECs under tri-culture showed 1.87-fold decrease in the rate of change in intensity over the initial intensity (ΔI/I_t=0_) when inhibitor was pre-treated. In detail for BBB triculture model, average ΔI/I_t=0_ was 0.7097 in DMSO control samples and 0.3798 in inhibitor treated samples. From this result, we infer that the p-gp inhibitor had a negative effect on the cellular system, allowing efflux calcein molecules to accumulate in the intracellular region. In contrast, EC monoculture samples did not exhibit a significant difference in the rate of change in intensity between inhibitor-treated (average ΔI/I_t=0_=0.8379) and DMSO control (average ΔI/I_t=0_=0.8376) samples (Fig. 4F). This result indicates that the p-gp inhibitor had little effect on ECs cultured in a non-BBB microenvironment.

### 3.5. Matrix modification enhanced astrocyte-EC interaction

The ECM plays a key role in regulating distinct cellular physiology in different tissues. Brain tissue surrounding the BBB is rich in ECM materials such as hyaluronic acid (HA), proteoglycans, and tenascins [35]. The critical role of HA in regulating the astrocyte phenotype *in vitro* has been reported previously [36–38].

To further recapitulate the distinct star-shaped morphology of astrocytes surrounding blood vessels, we tested the effect of HA in our system. We added HA hydrogel in different ratios, and then cultured HBMECs and astrocytes together until the vessel network formed in the central channel (Fig. 5A). As the proportion of HA was increased, astrocytes displayed a more star-shape morphology. When fibrin and HA were mixed at a 1:1 ratio (final concentrations of both fibrin and HA were 2.5 mg/mL), astrocytes were more branched than in fibrin gel only. A similar result was obtained when fibrin and HA were mixed at a ratio of 2.5:1.5 (final concentrations of fibrin and HA were 2.5 and 1.5 mg/mL, respectively), and most astrocytes had protrusions. This result agreed well with previous reports [36–38].

**Figure 5.**
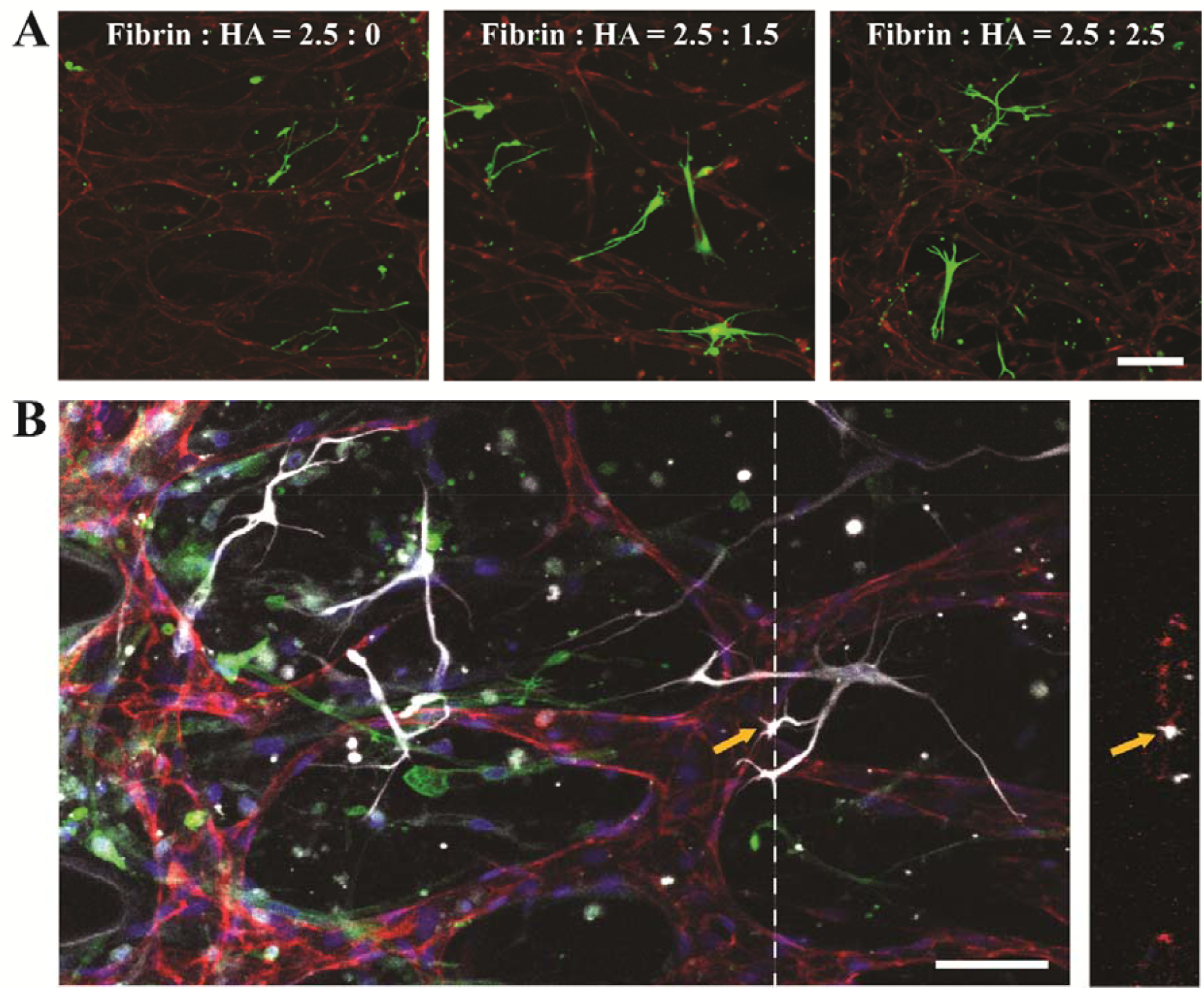
Matrix tuning by adding hyaluronic acid (HA) for improvement of astrocyte-EC interaction. (A) 3D confocal image of astrocytes (anti-GFAP, green) and HBMECs (anti-CD31, red) in each ECM hydrogel condition having different ratio of fibrin and HA mixture. Scale bar: 100μm. (B) Representative 3D confocal image of tri-culture of HBMECs (anti-CD31, red), astrocytes (anti-GFAP, white) and pericytes (anti-αSMA, green) in optimally tuned ECM, fibrin and HA mixed in 2.5:1.5 ratio. Nucleus were labeled by Hoeschst 33342 (blue). Right image is the section view of white dotted line in left image, describing direct contact of astrocyte end-feet on vascular surface (yellow arrow head). Scale bar: 70μm.

Vascular morphology, such as vessel width and lumen structure, was not significantly affected up to the addition of 1.5 mg/mL HA, whereas the effect of HA on astrocyte morphology was saturated at this ratio. Thus, we used a 2.5:1.5 mixture of fibrin and HA to perform a BBB tri-culture assay in the ECM. In the tri-culture model, astrocytes displayed a well-differentiated star-shaped morphology and more protrusions than the line-shaped astrocytes compare to fibrin-only hydrogel. These protrusions projected more astrocyte end-feet on endothelium, which may implicate increased astrocyte–EC interaction in our system (Fig. 5B).

## 4. Discussion

The development of an *in vitro* model capturing the complex physiology and barrier function of the BBB is crucial for understanding its biology, and for the design of CNS drugs. In this study, we reported a novel *in vitro* model that mimics BBB development via the mechanism of CNS angiogenesis, lending the model high physiological relevance. We demonstrated the importance of the 3D microenvironment in establishing the BBB phenotype, and the functionality.

Since *in vivo* BBB vasculature is distinctively developed through angiogenesis, and its functional phenotypes are emerged during the maturation process [4, 20] (Fig. 1B), our main strategy for attaining an *in vivo*-like model was to mimic the CNS angiogenesis. Our microfluidic model allowed long-term culture (> 1 week), and the vasculature had matured sufficiently to obtain BBB phenotypes. Morphological phenotypes that recapitulate BBB characters, such as limited vessel dilation (Fig. 2B and C), astrocyte end-foot and pericyte coverage on the vessels (Fig. E, Supplementary Video 1), were observed about a week of culture; these phenotypes were not visible in early stages of culture (Fig. 1D). Thus, mimicking the biological process of BBB development allowed the maturation of *in vivo*-like phenotypes in our model.

We showed critical role of cellular interaction among three cell types constituting BBB in our *in vitro* model (Fig. 2). BBB vessel is distinctly narrower than other tissue microvasculature [39, 40]. We clearly demonstrated that the diameter of vasculature was narrowest under tri-culture conditions (Fig. 2D). This trend highlights critical role of endothelial-perivascular cell communication on regulating vascular morphology in our model, although the diameter was slightly larger than known human brain capillaries (8–10μm) [40]. As we know, the average diameter of 34.64μm in this study is the smallest engineered vessel diameter that have been reported hitherto. The origin of ECs also seemed to be responsible for the specific BBB vascular phenotype (Supplementary Fig. 2). Using HUVECs originated from umbilical cord failed to represent the characteristic narrow-diameter BBB vasculature. Moreover, co-culture with different types of perivascular cells resulted different tendency compared to that of HBMECs. Junction expression was also distinguishable in our tri-culture system, unlike in the mono-culture system. We focused on the ZO-1 tight junction for each co-culture condition. Based on immunofluorescence analysis (Fig. 3A) and western blotting analysis (Fig. 3B), tri-culture conditions produced much clear and stronger ZO-1 expression than HBMEC monoculture conditions. All of these aforementioned analyses support the importance of using appropriate cell source and providing comprehensive cellular interactions in order to reconstitute the model which would further applied in drug development or disease studies targeted for human.

Our new 3D *in vitro* BBB system was shown to highly resemble the human BBB in terms of morphological phenotypes; by adopting the mechanism of CNS angiogenesis, we attempted to improve platform functionality based on these phenotypes. We focused on reconstituting the metabolic barrier function of the BBB, which is mainly responsible for preventing CNS drugs from crossing the endothelium into the brain region [29, 30, 41]. By taking advantage of our perfusable vascular network that allowed us to introduce particles or drugs from intraluminal side (Supplementary Fig. 1), we could flow calcein-AM molecules through the vessel network, which mimicked the circulation of drug molecules in the blood stream (Fig. 4B). As we tracked the decrease of fluorescence intensity over time (Fig. 4E), the efflux rate in BBB tri-culture system showed lower than that of EC mono-culture system. This result was different from our initial hypothesis that the efflux rate would be higher in BBB model due to increased expression of p-gp. However, transport system in BBB is not only limited to p-gp, but it is orchestrated by various modes such as carrier-mediated, receptor-mediated or paracellular transport [42–44]. This complex system could not be fully reconstituted in our *in vitro* model, therefore, this should be targeted to improve functional physiology in following future studies.

Nevertheless, our model prominently showed that the calcein molecule efflux rate was influenced by Valsopodar, a conventional inhibitor of multidrug resistant proteins such as p-gp, only when the vasculature was tri-cultured in the BBB microenvironment. In contrast, Valsopodar had no significant effect on 3D monocultured HBMECs (Fig. 4F). This result implicates that ECs in BBB tri-culture model possess p-gp as a dominant efflux transport system. Cellular interaction and BBB microenvironment during maturation process may have contributed in this phenotype. In short, the model showed potential as a novel screening platform for numerous p-gp inhibitors that will eventually contribute to improving the efficacy of CNS medications [30, 34]. Several assays and models have been used to study multi-drug-resistant proteins to modulate barrier function, however, none have attempted to reconstitute the *in vivo* microenvironment of BBB. Although the ultimate goal of regulating this protein is to increase the efficacy of human CNS drug delivery, most tests have been performed in mouse models [45, 46] and some human cell-based platforms have been restricted to 2D culture assays [33, 47, 48] with low physiological relevancy. Our platform, which allows for p-gp modulator testing in a highly *in* vivo-like microenvironment, will overcome these limitations and provide more reliable preclinical test results.

In 3D organ-on-a-chip model, ECM is the key component governing cell behaviors within the modeled tissues. Brain tissue is composed of complex ECM components, including HA, proteoglycans, and tenascins [35]. Among cells in the human BBB microenvironment, astrocyte phenotype and morphology are known to be largely influenced by the local microenvironment [36, 38]. This was clearly demonstrated by Placone et al. [37], who manipulated the type and ratio of ECM including HA, and analyzed astrocyte morphology and active states under each ECM condition. To recapitulate the distinct star-shaped astrocyte morphology in the 3D *in vitro* model, we tuned the 3D ECM base using previously developed protocols [37, 49]. In our microfluidic devices, astrocytes exhibited a star-shaped morphology in the presence of HA when mixed with fibrin hydrogel (Fig. 5A). By tuning the hydrogel, we were able to obtain astrocytes with many protrusions; this morphology was crucial for enhancing astrocyte-EC interactions, such as outer vessel wall end-foot structures (Fig. 5B). This is more than simply mimicking morphological structure since numerous studies have reported that astrocyte end-feet enhances the role of astrocytes in controlling homeostasis and repairing vascular damage in the CNS [3, 23, 50].

Our *in vitro* model was designed to represent the healthy BBB. However, many CNS diseases are intimately involved with the breakdown or dysfunction of the BBB. Obermeier et al. [2] reviewed the effect of the BBB on the pathology of diseases such as stroke, epilepsy, Alzheimer’s disease, and Parkinson’s disease. For example, dysfunction of the BBB efflux barrier system in clearing amyloid-β is known to be responsible for Alzheimer’s disease [51, 52], and the loss of pericytes in the BBB causes the pathology [53]. Since BBB cell types are selectively applicable, we expect our platform can provide useful tool to dissect the effects of each cell type on BBB pathology. Further development of our model to mimic BBB pathologies and to test therapies for them would be the next step in applying the model to translational research.

## 5. Conclusion

In this study, we reported a novel 3D *in vitro* BBB model that mimicked development of CNS angiogenesis. This strategy relies on biological knowledge that the maturation of BBB functional and morphological phenotypes occurs via angiogenesis, which is different from typical vasculature development of other tissues. In this platform, primary human brain ECs exhibited BBB phenotypes such as narrow vascular morphology, high ZO-1 expression, and a functional metabolic barrier system only when both perivascular cells, human astrocytes and pericytes, were present. This finding indicates that complex signaling and interaction through direct contact are indispensable for the reconstitution of an *in vitro* BBB microenvironment. Moreover, tuning the 3D hydrogel matrix by adding HA on fibrin-based hydrogel caused astrocytes to show their typical *in vivo* star-shaped structure, thereby enhancing interaction with the endothelium via end-foot structures.

We verified the functionality of the model by testing a BBB metabolic efflux transporter inhibitor, p-gp. Because p-gp modulation has been studied widely as an opportunity to enhance CNS pharmacotherapy, several inhibitors were screened prior to the study using *in vivo* models or simple 2D assays. Our experimental results showed that the efflux rates of calcein molecules as a substrate of p-gp were significantly decreased when the inhibitor was pretreated only in the BBB tri-culture system, but not in the EC monoculture. Based on this analysis, we assert that increasing the physiological relevance of the *in vitro* platform can contributes to the development of BBB morphological characteristics and functional phenotypes. Furthermore, the highly perfusable vasculature embedded in the 3D BBB microenvironment in our model can potentially be applied as a system for screening CNS drug candidates, as shown by the results of the p-gp inhibitor test. Combining this BBB tissue engineering with the emerging technology of high-throughput screening systems [54, 55] will permit the practical use of this model in pre-clinical tests and translational research, to develop improved CNS medications in the near future.

## 6. Disclosure

The authors report no conflicts of interest in this work.

## 7. Acknowledgement

This study was supported by Basic Science Research Program through the National Research Foundation of Korea (NRF) funded by the Ministry of Science and Technology (NRF-2016R1A4A1010796, 2018R1A2A1A05019550). In addition, it was supported by Global Ph.D. Fellowship Program also funded by the Ministry of Science and Technology (NRF-2016H1A2A1907378). Special thanks to Dr. Sujin Hyung and Seung-Ryeol Lee for helping western blot assay and Dr. Tae-Eun Park for providing experimental comments on efflux transport assay.

---

**Supplementary Figure 1A.**
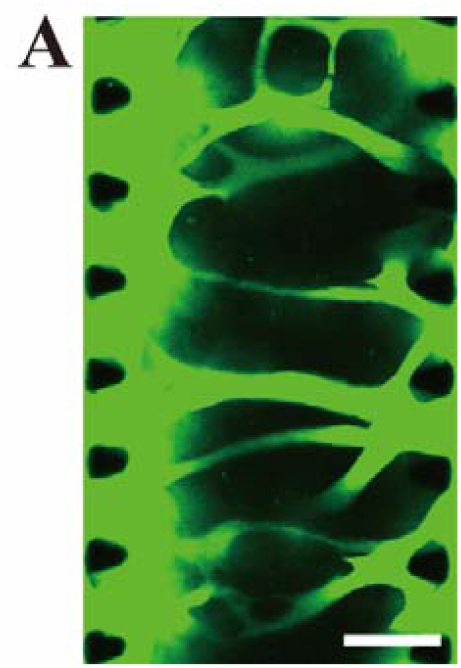
Tri-cultured 3D BBB *in vitro* microvasculature was perfused and each end was opened. This was verified by 3D confocal imaging after flowing FITC-Dextran (70kDa) from left side to right end. Scale bar: 200 μm.

**Supplementary Figure 4.**
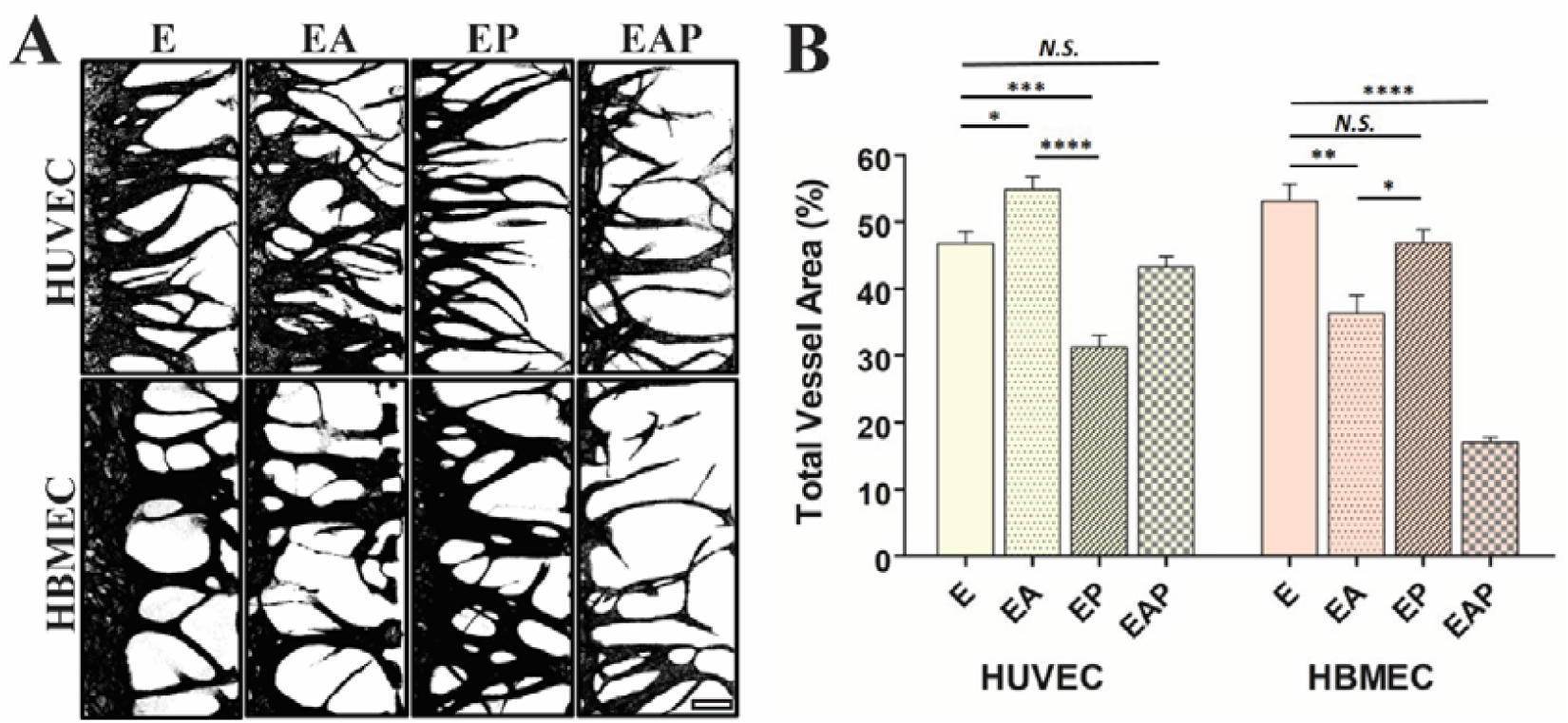
Analysis of vascular morphology depending on origin of endothelium. (A) Masked images of 3D vasculature in each co-culture condition using either HUVEC or HBMEC, fixed at day 5 or day 7, respectively. Scale bar: 200μm. (B) Quantitative analysis on total vasculature in each co-culture condition. For HUVEC, n=8 in condition E and EP; n=6 in condition EA; n=12 in condition EAP. For HBMEC, n=8 in condition E, EA and EP; n=10 in condition EAP. Error bars represent SEM. Unpaired two-tailed Student’s t-test was performed to obtain statistical comparisons of analyzed values, with the *p* value threshold for statistical significance set at *p<0.05; **p<0.005; ***p<0.0005; ****p<0.00001.

**Supplementary Figure 3A.**
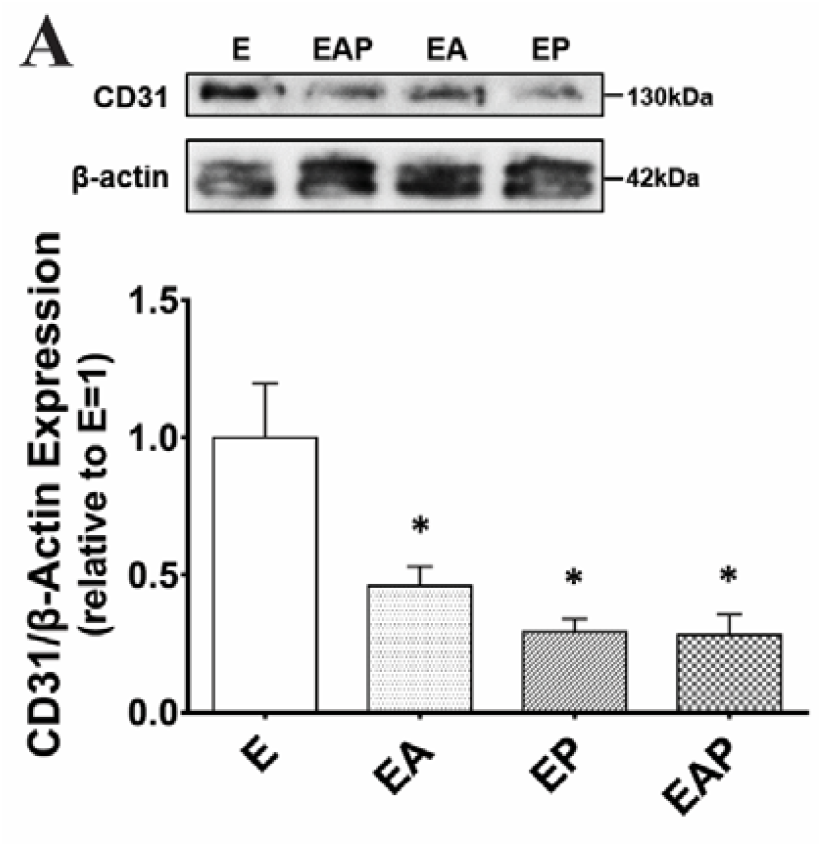
Western blot assay for quantifying CD31 expression. Upper image is representative band image of CD31 and β-actin for each co-culture condition. Below graph is the result of quantitative analysis on CD31 expression level in each co-culture condition normalized against the expression level of β-actin. N=6 for condition E, EA and EAP; n=5 for condition EP. Error bars represent SEM. Unpaired two-tailed Student’s t-test was performed to obtain statistical comparisons of analyzed values, with the *p* value threshold for statistical significance set at *p<0.05.

**Supplementary Video 1.**
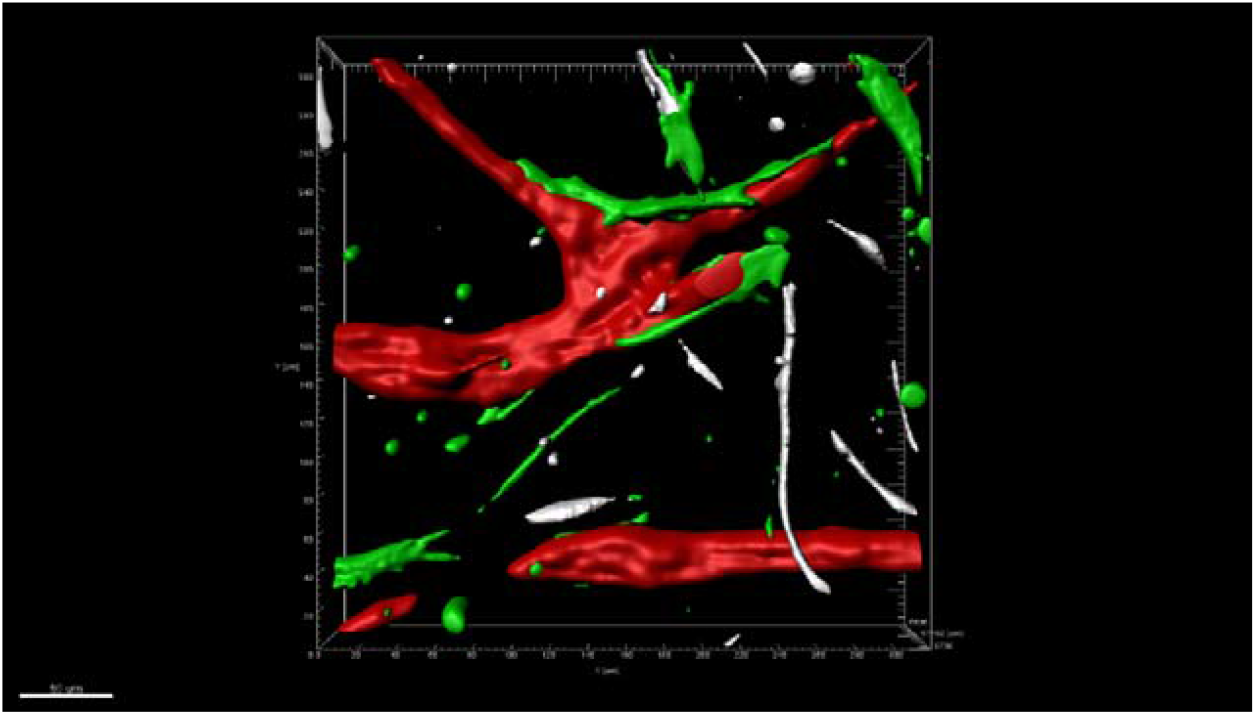
Software reconstruction of 3D BBB tri-cultured model image (Figure 1E) from confocal microscopy which clearly demonstrates cellular interaction between cells. 3D lumenized HBMEC (red) is covered by pericytes (green) and astrocytes (white) possess end-feet on surface of microvessel.

## References

[1] W.A. Banks, From blood-brain barrier to blood-brain interface: new opportunities for CNS drug delivery, Nat Rev Drug Discov 15(4) (2016) 275–92.

[2] B. Obermeier, R. Daneman, R.M. Ransohoff, Development, maintenance and disruption of the blood-brain barrier, Nat Med 19(12) (2013) 1584–96.

[3] N.J. Abbott, L. Ronnback, E. Hansson, Astrocyte-endothelial interactions at the blood-brain barrier, Nat Rev Neurosci 7(1) (2006) 41–53.

[4] B. Engelhardt, Development of the blood-brain barrier, Cell Tissue Res 314(1) (2003) 119–29.

[5] M. Vallon, J. Chang, H. Zhang, C.J. Kuo, Developmental and pathological angiogenesis in the central nervous system, Cell Mol Life Sci 71(18) (2014) 3489–506.

[6] R.A. Umans, H.E. Henson, F. Mu, C. Parupalli, B. Ju, J.L. Peters, K.A. Lanham, J.S. Plavicki, M.R. Taylor, CNS angiogenesis and barriergenesis occur simultaneously, Dev Biol 425(2) (2017) 101–108.

[7] S. Nakagawa, M.A. Deli, H. Kawaguchi, T. Shimizudani, T. Shimono, A. Kittel, K. Tanaka, M. Niwa, A new blood-brain barrier model using primary rat brain endothelial cells, pericytes and astrocytes, Neurochem Int 54(3–4) (2009) 253–63.

[8] S. Nakagawa, M.A. Deli, S. Nakao, M. Honda, K. Hayashi, R. Nakaoke, Y. Kataoka, M. Niwa, Pericytes from brain microvessels strengthen the barrier integrity in primary cultures of rat brain endothelial cells, Cell Mol Neurobiol 27(6) (2007) 687–94.

[9] K. Hatherell, P.O. Couraud, I.A. Romero, B. Weksler, G.J. Pilkington, Development of a three-dimensional, all-human in vitro model of the blood-brain barrier using mono-, co-, and tri-cultivation Transwell models, J Neurosci Methods 199(2) (2011) 223–9.

[10] R. Booth, H. Kim, Characterization of a microfluidic in vitro model of the blood-brain barrier (muBBB), Lab Chip 12(10) (2012) 1784–92.

[11] A.K. Achyuta, A.J. Conway, R.B. Crouse, E.C. Bannister, R.N. Lee, C.P. Katnik, A.A. Behensky, J. Cuevas, S.S. Sundaram, A modular approach to create a neurovascular unit-on-a-chip, Lab Chip 13(4) (2013) 542–53.

[12] J.A. Brown, V. Pensabene, D.A. Markov, V. Allwardt, M.D. Neely, M. Shi, C.M. Britt, O.S. Hoilett, Q. Yang, B.M. Brewer, P.C. Samson, L.J. McCawley, J.M. May, D.J. Webb, D. Li, A.B. Bowman, R.S. Reiserer, J.P. Wikswo, Recreating blood-brain barrier physiology and structure on chip: A novel neurovascular microfluidic bioreactor, Biomicrofluidics 9(5) (2015) 054124.

[13] L.M. Griep, F. Wolbers, B. de Wagenaar, P.M. ter Braak, B.B. Weksler, I.A. Romero, P.O. Couraud, I. Vermes, A.D. van der Meer, A. van den Berg, BBB on chip: microfluidic platform to mechanically and biochemically modulate blood-brain barrier function, Biomed Microdevices 15(1) (2013) 145–50.

[14] K.L. Sellgren, B.T. Hawkins, S. Grego, An optically transparent membrane supports shear stress studies in a three-dimensional microfluidic neurovascular unit model, Biomicrofluidics 9(6) (2015) 061102.

[15] J.D. Wang, S. Khafagy el, K. Khanafer, S. Takayama, M.E. ElSayed, Organization of Endothelial Cells, Pericytes, and Astrocytes into a 3D Microfluidic in Vitro Model of the Blood-Brain Barrier, Mol Pharm 13(3) (2016) 895–906.

[16] H. Cho, J.H. Seo, K.H. Wong, Y. Terasaki, J. Park, K. Bong, K. Arai, E.H. Lo, D. Irimia, Three-Dimensional Blood-Brain Barrier Model for in vitro Studies of Neurovascular Pathology, Sci Rep 5 (2015) 15222.

[17] A. Herland, A.D. van der Meer, E.A. FitzGerald, T.E. Park, J.J. Sleeboom, D.E. Ingber, Distinct Contributions of Astrocytes and Pericytes to Neuroinflammation Identified in a 3D Human Blood-Brain Barrier on a Chip, PLoS One 11(3) (2016) e0150360.

[18] G. Adriani, D. Ma, A. Pavesi, R.D. Kamm, E.L. Goh, A 3D neurovascular microfluidic model consisting of neurons, astrocytes and cerebral endothelial cells as a blood-brain barrier, Lab Chip 17(3) (2017) 448–459.

[19] S. Bang, S.R. Lee, J. Ko, K. Son, D. Tahk, J. Ahn, C. Im, N.L. Jeon, A Low Permeability Microfluidic Blood-Brain Barrier Platform with Direct Contact between Perfusable Vascular Network and Astrocytes, Sci Rep 7(1) (2017) 8083.

[20] S. Liebner, C.J. Czupalla, H. Wolburg, Current concepts of blood-brain barrier development, Int J Dev Biol 55(4–5) (2011) 467–76.

[21] R. Daneman, D. Agalliu, L. Zhou, F. Kuhnert, C.J. Kuo, B.A. Barres, Wnt/beta-catenin signaling is required for CNS, but not non-CNS, angiogenesis, Proc Natl Acad Sci U S A 106(2) (2009) 641–6.

[22] S. Kim, H. Lee, M. Chung, N.L. Jeon, Engineering of functional, perfusable 3D microvascular networks on a chip, Lab Chip 13(8) (2013) 1489–500.

[23] L.G. Dubois, L. Campanati, C. Righy, I. D’Andrea-Meira, T.C. Spohr, I. Porto-Carreiro, C.M. Pereira, J. Balca-Silva, S.A. Kahn, M.F. DosSantos, A. Oliveira Mde, A. Ximenes-da-Silva, M.C. Lopes, E. Faveret, E.L. Gasparetto, V. Moura-Neto, Gliomas and the vascular fragility of the blood brain barrier, Front Cell Neurosci 8 (2014) 418.

[24] Y. Yao, Z.L. Chen, E.H. Norris, S. Strickland, Astrocytic laminin regulates pericyte differentiation and maintains blood brain barrier integrity, Nat Commun 5 (2014) 3413.

[25] A. ElAli, P. Theriault, S. Rivest, The role of pericytes in neurovascular unit remodeling in brain disorders, Int J Mol Sci 15(4) (2014) 6453–74.

[26] N.B. Hamilton, D. Attwell, C.N. Hall, Pericyte-mediated regulation of capillary diameter: a component of neurovascular coupling in health and disease, Front Neuroenergetics 2 (2010).

[27] A. Zozulya, C. Weidenfeller, H.J. Galla, Pericyte-endothelial cell interaction increases MMP-9 secretion at the blood-brain barrier in vitro, Brain Res 1189 (2008) 1–11.

[28] A. Armulik, G. Genove, M. Mae, M.H. Nisancioglu, E. Wallgard, C. Niaudet, L. He, J. Norlin, P. Lindblom, K. Strittmatter, B.R. Johansson, C. Betsholtz, Pericytes regulate the blood-brain barrier, Nature 468(7323) (2010) 557–61.

[29] H. Kusuhara, H. Suzuki, T. Terasaki, A. Kakee, M. Lemaire, Y. Sugiyama, P-Glycoprotein Mediates the Efflux of Quinidine across the Blood-Brain Barrier, Journal of Pharmacology and Experimental Therapeutics 283(2) (1997) 574–580.

[30] D.S. Miller, B. Bauer, A.M. Hartz, Modulation of P-glycoprotein at the blood-brain barrier: opportunities to improve central nervous system pharmacotherapy, Pharmacol Rev 60(2) (2008) 196–209.

[31] A.H. Schinkel, P-Glycoprotein, a gatekeeper in the blood–brain barrier, Advanced Drug Delivery Reviews 36(2-3) (1999) 179–194.

[32] A. Tsuji, I. Tamai, Blood-brain barrier function of P-glycoprotein, Advanced Drug Delivery Reviews 25(2–3) (1997) 287–298.

[33] Z. Holló, L. Homolya, C.W. Davis, B. Sarkadi, Calcein accumulation as a fluorometric functional assay of the multidrug transporter, Biochimica et Biophysica Acta (BBA) - Biomembranes 1191(2) (1994) 384–388.

[34] F. Tiberghien, F. Loor, Ranking of P-glycoprotein substrates and inhibitors by a calcein-AM fluorometry screening assay, Anticancer Drugs 7(5) (1996) 568–578.

[35] D.R. Zimmermann, M.T. Dours-Zimmermann, Extracellular matrix of the central nervous system: from neglect to challenge, Histochem Cell Biol 130(4) (2008) 635–53.

[36] J. Arulmoli, H.J. Wright, D.T.T. Phan, U. Sheth, R.A. Que, G.A. Botten, M. Keating, E.L. Botvinick, M.M. Pathak, T.I. Zarembinski, D.S. Yanni, O.V. Razorenova, C.C.W. Hughes, L.A. Flanagan, Combination scaffolds of salmon fibrin, hyaluronic acid, and laminin for human neural stem cell and vascular tissue engineering, Acta Biomater 43 (2016) 122–138.

[37] A.L. Placone, P.M. McGuiggan, D.E. Bergles, H. Guerrero-Cazares, A. Quinones-Hinojosa, P.C. Searson, Human astrocytes develop physiological morphology and remain quiescent in a novel 3D matrix, Biomaterials 42 (2015) 134–43.

[38] A.F. Levy, M. Zayats, H. Guerrero-Cazares, A. Quinones-Hinojosa, P.C. Searson, Influence of basement membrane proteins and endothelial cell-derived factors on the morphology of human fetal-derived astrocytes in 2D, PLoS One 9(3) (2014) e92165.

[39] A. Mishra, F.M. O’Farrell, C. Reynell, N.B. Hamilton, C.N. Hall, D. Attwell, Imaging pericytes and capillary diameter in brain slices and isolated retinae, Nat Protoc 9(2) (2014) 323–36.

[40] A.D. Wong, M. Ye, A.F. Levy, J.D. Rothstein, D.E. Bergles, P.C. Searson, The blood-brain barrier: an engineering perspective, Front Neuroeng 6 (2013) 7.

[41] A.G. de Boer, I.C. van der Sandt, P.J. Gaillard, The role of drug transporters at the blood-brain barrier, Annu Rev Pharmacol Toxicol 43 (2003) 629–56.

[42] C. Hajal, M. Campisi, C. Mattu, V. Chiono, R.D. Kamm, In vitro models of molecular and nano-particle transport across the blood-brain barrier, Biomicrofluidics 12(4) (2018) 042213.

[43] V. Neves, F. Aires-da-Silva, S. Corte-Real, M.A.R.B. Castanho, Antibody Approaches To Treat Brain Diseases, Trends in Biotechnology 34(1) (2016) 36–48.

[44] R. Gabathuler, Approaches to transport therapeutic drugs across the blood-brain barrier to treat brain diseases, Neurobiol Dis 37(1) (2010) 48–57.

[45] S. Bihorel, G. Camenisch, M. Lemaire, J.M. Scherrmann, Influence of breast cancer resistance protein (Abcg2) and p-glycoprotein (Abcb1a) on the transport of imatinib mesylate (Gleevec) across the mouse blood-brain barrier, J Neurochem 102(6) (2007) 1749–1757.

[46] A.H. Schinkel, J.J.M. Smit, O. van Tellingen, J.H. Beijnen, E. Wagenaar, L. van Deemter, C.A.A.M. Mol, M.A. van der Valk, E.C. Robanus-Maandag, H.P.J. te Riele, A.J.M. Berns, P. Borst, Disruption of the mouse mdr1a P-glycoprotein gene leads to a deficiency in the blood-brain barrier and to increased sensitivity to drugs, Cell 77(4) (1994) 491–502.

[47] D.G. Kim, M.S. Bynoe, A2A adenosine receptor modulates drug efflux transporter P-glycoprotein at the blood-brain barrier, J Clin Invest 126(5) (2016) 1717–33.

[48] P.J. Gaillard, I.C.J. van der Sandt, L.H. Voorwinden, D. Vu, J.L. Nielsen, A.G. de Boer, D.D. Breimer, Astrocytes Increase the Functional Expression of PGlycoprotein in an In Vitro Model of The Blood-Brain Barrier, Pharmaceutical Research 17(10) (2000) 1198–1205.

[49] P.P. Partyka, G.A. Godsey, J.R. Galie, M.C. Kosciuk, N.K. Acharya, R.G. Nagele, P.A. Galie, Mechanical stress regulates transport in a compliant 3D model of the blood-brain barrier, Biomaterials 115 (2017) 30–39.

[50] R.F. Haseloff, I.E. Blasig, H.C. Bauer, H. Bauer, In Search of the Astrocytic Factor(s) Modulating Blood–Brain Barrier Functions in Brain Capillary Endothelial Cells In Vitro, Cellular and Molecular Neurobiology 25(1) (2005) 25–39.

[51] J.M. Tarasoff-Conway, R.O. Carare, R.S. Osorio, L. Glodzik, T. Butler, E. Fieremans, L. Axel, H. Rusinek, C. Nicholson, B.V. Zlokovic, B. Frangione, K. Blennow, J. Menard, H. Zetterberg, T. Wisniewski, M.J. de Leon, Clearance systems in the brain-implications for Alzheimer disease, Nat Rev Neurol 11(8) (2015) 457–70.

[52] F. Marques, J.C. Sousa, N. Sousa, J.A. Palha, Blood-brain-barriers in aging and in Alzheimer’s disease, Mol Neurodegener 8 (2013) 38.

[53] A.P. Sagare, R.D. Bell, Z. Zhao, Q. Ma, E.A. Winkler, A. Ramanathan, B.V. Zlokovic, Pericyte loss influences Alzheimer-like neurodegeneration in mice, Nat Commun 4 (2013) 2932.

[54] D. Barata, C. van Blitterswijk, P. Habibovic, High-throughput screening approaches and combinatorial development of biomaterials using microfluidics, Acta Biomater 34 (2016) 1–20.

[55] S. Lee, J. Ko, D. Park, S.R. Lee, M. Chung, Y. Lee, N.L. Jeon, Microfluidic-based vascularized microphysiological systems, Lab Chip (2018).

